# Coordinated Representational Drift Across the Mouse Cortex

**DOI:** 10.64898/2026.05.05.723038

**Authors:** Ryan Peters, James Hope, Michael Feldkamp, Travis Beckerle, Ibrahim Oladepo, Ihor Hryb, Kapil Saxena, A. David Redish, Suhasa Kodandaramaiah

## Abstract

Cortical neurons continuously change their spatial tuning properties over days and weeks, yet how this representational drift is coordinated across distributed cortical networks remains unknown. Using a robotic cranial exoskeleton for chronic widefield calcium imaging with cellular resolution, we tracked over 110,000 unique layer 2/3 neurons across retrosplenial, visual, somatosensory, and motor cortices in mice navigating a figure-8 maze spanning 47 days. Single-neuron spatial tuning properties followed a posterior-to-anterior gradient, with retrosplenial and visual cortices containing the highest proportions of spatially tuned neurons. This was dissociated from local functional coupling, which was strongest in somatosensory cortex. Population activity formed a low-dimensional manifold whose geometry mirrored the structure of the maze. Despite these regional differences, all four regions decorrelated with similar exponential timescales, and session-specific deviations from each region’s decay trajectory were correlated across all pairwise region combinations, suggesting that drift was coordinated rather than independent across the dorsal cortex. This coordination persisted after controlling for shared behavioral fluctuations. Furthermore, drift was consistent with an orthogonal transformation of the population code that preserved the geometric relationships between spatial representations across sessions despite continuous drift in single-neuron tuning.

## Introduction

Individual neurons in the mammalian cortex continuously change their tuning properties across days and weeks even as animals perform familiar tasks with stable behavior in unchanging environments, a phenomenon termed representational drift [9, 32]. Representational drift has been documented across a wide range of brain areas, including primary sensory cortices [22, 7, 3] and the hippocampus [20, 13, 19], raising a fundamental question about how neural systems maintain stable computation despite continuously changing neural representations. Current studies have largely examined drift within individual brain areas, leaving open whether representational drift operates through mechanisms that are local and independent within each cortical region or are coordinated across distributed networks.

Spatial navigation provides a particularly well-suited framework to study representational drift across time [32, 27], as it engages multiple cortical regions simultaneously, including retrosplenial, visual, motor, and somatosensory cortices. Prior work has shown that within individual regions, drift is consistent with orthogonal transformations of the population code between sessions, which preserve the geometric relationships between neural states even as individual neurons change their tuning [9, 12]. If such transformations are globally coordinated, rather than occurring independently within each region, this would imply a unified organizational principle maintaining representational stability across the cortex as a whole.

However, chronic and simultaneous recording from large populations of neurons across multiple cortical areas in freely behaving animals has proven technically challenging. Previous studies have faced fundamental trade-offs between spatial coverage, resolution, and the ability to record from freely moving animals over extended timescales. Head-fixed preparations and virtual reality environments [15, 35, 34, 2, 28, 4, 23] enable stable imaging with high resolution, but alter the sensorimotor experience of navigation in ways that may affect cortical representations.

We addressed these limitations using a robotic cranial exoskeleton [17] that enabled chronic widefield single-photon calcium imaging with cellular resolution across the dorsal cortex in mice navigating a physical figure-8 maze. Using the cranial exoskeleton, we tracked over 110,000 unique layer 2/3 neurons across retrosplenial, visual, somatosensory, and motor cortices spanning 47 days. We first characterized single-neuron spatial tuning properties across regions, revealing a posterior-to-anterior gradient in spatial coding that was dissociated from local functional coupling. We then examined the temporal evolution of these representations at the population level. All four regions decorrelated with similar exponential timescales, and session-specific deviations from each region’s decay trajectory were correlated across all region pairs, demonstrating that drift was coordinated rather than independent across the dorsal cortex. By leveraging the simultaneously recorded behavioral signals from the exoskeleton (position, velocity, acceleration, and applied force), we further showed that this drift could not be explained by changes in the animal’s behavior across sessions. Finally, geometric analysis revealed that this coordinated drift was consistent with orthogonal transformations of the population code, preserving the relative relationships between spatial representations across sessions despite continuous drift in single-neuron tuning. Together, these findings suggest a unified organizational principle in which representational stability across layer 2/3 cortex is maintained through population-level geometric invariance, even in the absence of stable individual neuron tuning.

## Results

### A robotic cranial exoskeleton enables large-scale cortical imaging in mice navigating a figure-8 maze

We performed cellular-resolution widefield single-photon calcium imaging across the dorsal cortex in mice navigating a figure-8 maze while docked to a robotically actuated imaging head-stage (**Fig. 1A,B**). The imaging headstage incorporated force sensors within the docking clamp that provided feedback to actuate it via a delta robot in response to the applied cranial forces along a pre-determined trajectory in the figure-8 maze (**Fig. 1A**; Methods A.2) [17]. The figure-8 maze contained two reward zones positioned on the left and right sides, with visual cues placed in the upper sections of each annulus (**Fig. 1A**).

**Figure 1:**
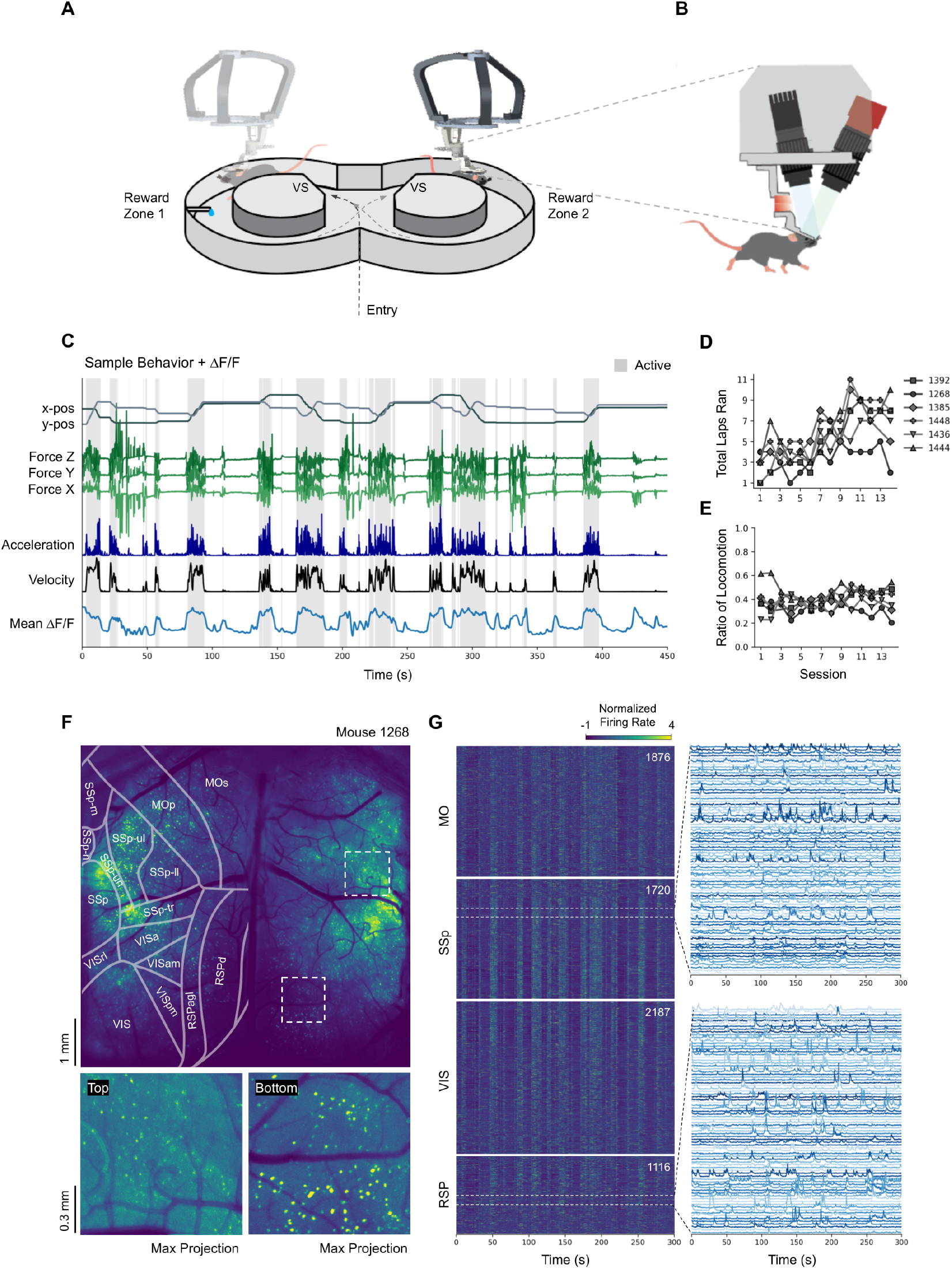
Widefield cellular-resolution imaging during navigation in a robotically actuated exoskeleton. **(A)** Schematic of the cranial exoskeleton imaging system within the figure-8 maze environment. Arrows show mouse trajectory within the maze. The locations of the visual stimuli are labeled as VS. **(B)** Magnified view of the imaging headstage. **(C)** Representative behavioral variables and mean ΔF/F. **(D)** Total laps completed per session for each mouse. **(E)** Proportion of time per session spent actively locomoting versus stationary. **(F)** Top: representative maximum intensity projection from Suite2p, registered to the Allen Brain Atlas. Bottom: magnified views from dorsal cortex in the right hemisphere. **(G)** Left: Normalized ΔF/F activity from mouse 1268, with cells grouped by cortical region and sorted within each region by anteroposterior position (most anterior at top). Per-region cell counts are shown in the top-right corner. Right: example ΔF/F traces from primary somatosensory cortex (SSp) and retrosplenial cortex (RSP).

Each session began with mice positioned on a circular treadmill for initial calibration before entering the maze environment. Mice then followed a predefined trajectory that the robotically actuated headstage could follow, with the system designed to guide forward motion and ensure consistent stopping at each maze terminus regardless of reward location. Upon stopping, a chocolate milk reward was automatically dispensed at active reward zones. This controlled movement protocol enabled circularization of the 2D spatial environment. The reward was dispensed on the left side during sessions 1–10 and on the right side during sessions 11–14. Each mouse performed between 13 and 14 sessions over the course of 47 days (**Fig. 1D**), with mice spending an average of 38.9 ± 8.3% (SD across sessions) of time actively locomoting in each session (**Fig. 1E**).

The robotically actuated headstage incorporated a single-photon microscope capable of imaging an 8.6 mm × 6.6 mm field of view at ∼3.45 µm pixel resolution in mice sparsely expressing GCaMP6s in layer 2/3 pyramidal neurons [8] (*n* = 6 mice, Cux2Cre-ERT2 [10] × Ai163 [6]) [18]. Cell extraction was performed using Suite2p [26], followed by manual validation and filtering based on spatiotemporal properties of the extracted cell footprints and ΔF/F. A representative maximum intensity projection image of one mouse during one imaging session is shown in **Fig. 1F** (all six mice in **Supplementary Fig. 1A**). Single-point tracking of the mouse position was obtained from the mechanical positioning of the exoskeleton itself, along with velocity, acceleration, and force sensor data for behavioral analysis (**Fig. 1C,D,E**). Consistent with previous studies, the onset of locomotion resulted in a broad increase in neural activity across the cortex (**Fig. 1C**). In total, over 110,000 unique neurons were extracted across mice (*n* = 6). Single neurons were simultaneously imaged bilaterally in the visual, somatosensory, retrosplenial, and motor cortices (**Fig. 1F,G**; per-mouse anatomical distribution in **Supplementary Fig. 1B**).

### Tracking cells in mesoscale calcium imaging across 47 days

To analyze longitudinal dynamics of individual neurons, we developed a cell registration algorithm to track neurons across recording sessions (**Fig. 2**). Due to the large number of detected cells and their broad spatial distribution across the dorsal cortex, we combined an optimized anatomical feature matching algorithm with an adapted Bayesian likelihood scoring method for cell registration (Methods A.6). Briefly, for each session, mean fluorescence projections and cell footprints were extracted from Suite2p and used for session alignment (**Fig. 2C,D**). A representative example of cells detected in two non-contiguous areas of the cortex is shown in **Fig. 2A**. Example traces of neuronal activity for some cells indicated in **Fig. 2A** are shown in **Fig. 2B**. The registration algorithm successfully tracked an average of ∼50% of the cells across any pair of sessions (**Fig. 2E**). The proportion of cells successfully tracked across *N* sessions closely followed a 1*/N* scaling (R^2^ = 0.957 ± 0.025 SD across mice), with approxi-mately 100*/N* % of cells being able to be tracked across *N* sessions. Furthermore, we found this scaling to be invariant to which sessions we chose to track between (**Fig. 2E,F**). Across all mice, we recorded between 763 and 8,300 neurons in each session. However, not all neurons imaged in a given session were active in subsequent sessions, with a subset of the neurons being active across multiple sessions. Pooling the unique neurons identified across all sessions, we tracked between 9,799 and 27,378 cells per mouse across all sessions (**Fig. 2G**).

**Figure 2:**
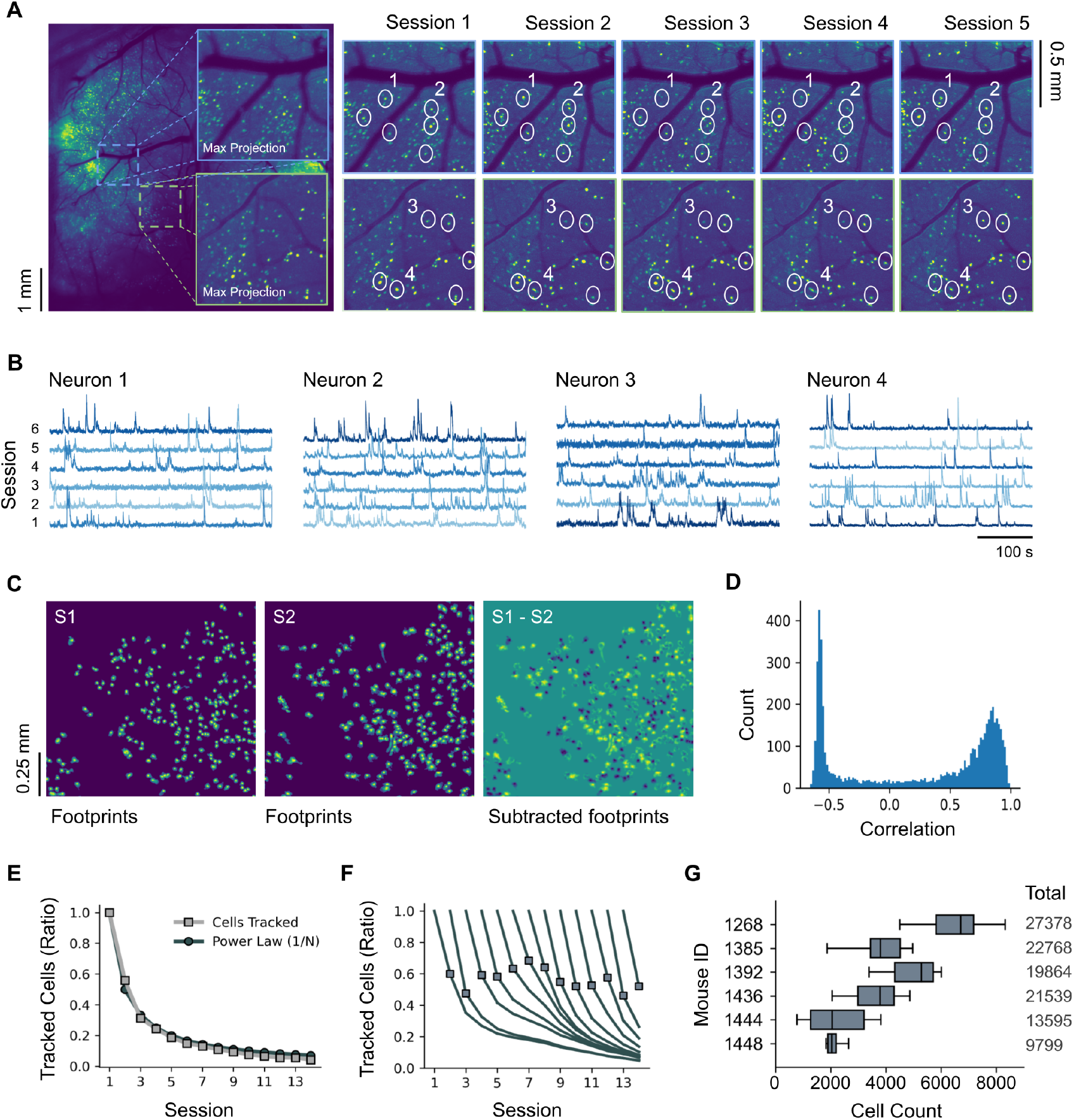
Longitudinal cell tracking across imaging sessions. **(A)** Representative cells tracked across 5 sessions. Maximum intensity projection from mouse 1268 (left) with magnified view of sessions 1–5 (right). Circles denote sample cells successfully tracked across all sessions. **(B)** Sample ΔF/F traces from the cells shown in (A), extended to 6 sessions. **(C)** Raw footprints extracted via Suite2p from session 1 (left) and session 2 (middle) from mouse 1268, with the pixelwise subtraction of the footprints after alignment (right). **(D)** Distribution of spatial correlations between registered cell footprints from sessions 1 and 2 shown in (C). **(E)** Proportion of cells successfully tracked across N consecutive sessions, averaged across mice. Shown alongside a 1/N power law (black line). **(F)** Proportion of cells tracked across N consecutive sessions with different starting sessions. Square markers label the number of cells tracked across two sessions. **(G)** Horizontal bars show per-session cell counts per mouse. Right-side tick labels indicate the total unique cells identified per mouse after registration.

### Single-cell spatial tuning across the cortex during active navigation

To characterize how individual neurons encode spatial information during navigation, we computed lapwise spatial tuning functions (LSTFs) for each neuron across all recording sessions. The figure-8 maze was divided into 36 equally sized spatial bins (∼4.45 cm each), and for each neuron we calculated per-lap tuning functions by averaging ΔF/F values within each bin. We excluded periods of immobility (*<*0.39 cm/s) to focus our analysis on active navigation epochs. ΔF/F traces were smoothed with a Gaussian filter (*σ* = 0.75 seconds) prior to binning, and tuning functions were z-score normalized within each session to account for baseline shifts in neural activity across sessions. LSTFs were organized as a three-dimensional tensor (laps × spatial bins × neurons) (**Fig. 3A**; Methods A.7).

**Figure 3:**
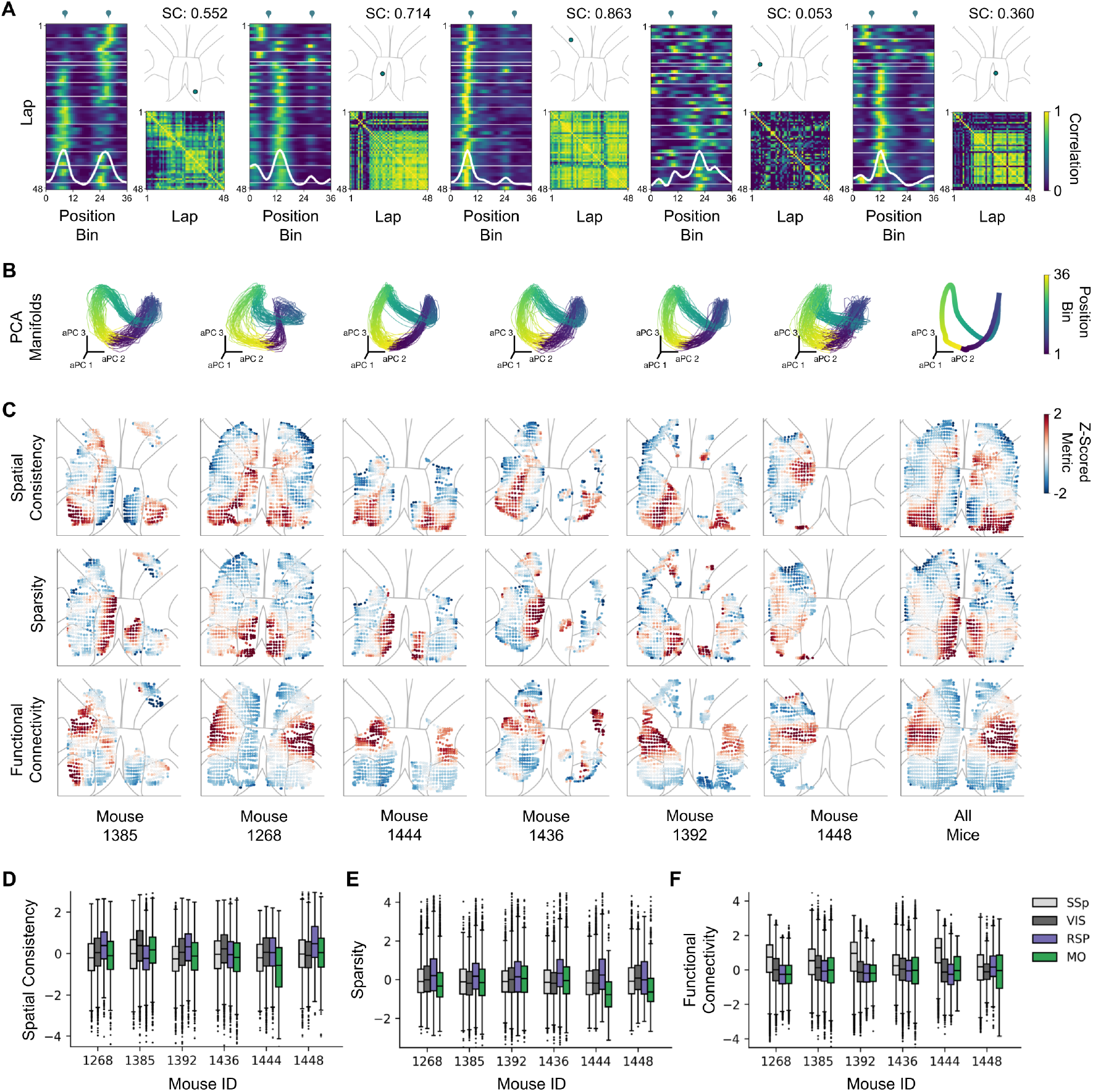
Spatial tuning properties and population dynamics across dorsal cortex. **(A)** Lapwise tuning functions from five representative neurons tracked across all sessions. Left: Position-by-lap tuning functions with horizontal white lines indicating session boundaries and droplet symbols marking reward zone locations. Top right: Anatomical location along dorsal cortex. Bottom right: Lap-by-lap cross-correlation matrix of the tuning function, with spatial consistency scores displayed in the upper right corner. **(B)** Population activity embedded in the first three aligned principal components (aPC1–3) for each mouse. Each point represents a spatial position bin on a given lap, colored by position along the track. PCs were aligned across mice by fitting a linear map to a reference mouse for visualization purposes. The rightmost panel shows the mean embedding averaged across all mice, laps, and sessions (Methods A.8). **(C)** Spatial distribution of tuning metrics across dorsal cortex. Metrics were computed within evenly spaced grid tiles and z-score normalized within each mouse. Top: spatial consistency. Middle: sparsity. Bottom: functional connectivity. **(D–F)** Spatial consistency (D), sparsity (E), and functional connectivity (F) values for each mouse, z-score normalized within each mouse.

Consistent with previous work [24, 38], single-neuron LSTFs exhibited a continuum of spatial selectivity, ranging from neurons with highly reliable spatial tuning to those with no discernible spatial preference (**Fig. 3A**). To quantify this variability, we computed a spatial consistency (SC) score for each neuron (Methods A.9). The anatomical distribution of SC scores across all neurons recorded in the dorsal cortex is shown in **Fig. 3C** (top row), with region-specific distributions shown in **Fig. 3D**.

Pooling all detected cells across the four cortical regions (*n* = 114,802), spatial consistency varied substantially between regions (pooled raw SC: 0.361 ± 0.235 SD across all mice and regions; **Fig. 3D**). After within-mouse z-scoring to remove baseline differences (Methods A.18), both posterior regions, visual cortex (VIS: +0.101 ± 0.005 SEM) and retrosplenial cortex (RSP: −0.007 ± 0.006 SEM), exhibited higher spatial consistency than the anterior regions (motor cortex, MO: −0.070 ± 0.007 SEM; primary somatosensory cortex, SSp: −0.143 ± 0.007 SEM). A linear mixed-effects model with mouse identity as a random intercept confirmed significant differences between regions, with post-hoc Tukey’s HSD pairwise comparisons all yielding *P <* 0.001 and the largest effect observed between VIS and SSp (Cohen’s *d* = 0.245). The proportion of neurons exceeding a spatial consistency threshold of 0.5 followed a similar posterior-to-anterior gradient (RSP: 34.4% ± 15.8% SD across mice; VIS: 32.6% ± 10.2%; MO: 24.3% ± 4.7%; SSp: 24.1% ± 8.9%), with VIS and RSP not differing significantly from each other (paired *t*-test, *P* = 0.788). RSP additionally exhibited pronounced bimodality across mice, with four animals at mean SC = 0.41–0.51 and two at 0.24–0.31 (before z-scoring; **Fig. 3C**, top row).

We next examined the spatial sparsity of neural responses (**Fig. 3C**, middle row). For each neuron, we computed a modified lifetime sparsity index from its spatial tuning function in each session and averaged across sessions [30, 40], where values near 1 indicate a neuron is actively firing in only a small number of spatial bins, while values near 0 indicate the neuron fires uniformly across all position bins (Methods A.10). Across the same set of neurons (z-scored as above), RSP exhibited the highest spatial sparsity (0.252 ± 0.007 SEM), significantly exceeding all other regions (all pairwise comparisons *P <* 0.001; largest effect: RSP vs. MO, Cohen’s *d* = 0.367; **Fig. 3E**). In contrast, VIS, SSp, and MO differed only modestly from one another (VIS: −0.042 ± 0.004 SEM; SSp: −0.066 ± 0.006 SEM; MO: −0.139 ± 0.007 SEM; all pairwise |*d*| *<* 0.14). This suggests that while both RSP and VIS contained neurons with reliable spatial tuning (as measured by SC), a higher fraction of RSP neurons were distinguished by particularly sharp spatial fields confined to fewer maze locations.

Finally, we characterized the functional connectivity of each recorded neuron, defined as the strength of distance-dependent correlation structure within a hemisphere (**Fig. 3C**, bottom row). For each neuron, we computed the Pearson correlation between its pairwise physical distances to all other neurons in the same hemisphere and the corresponding pairwise signal correlations, and then negated this value and clipped at zero so that positive functional connectivity scores indicated stronger local correlation structure (Methods A.11). Across all neurons (within-mouse z-scored), SSp showed the highest functional connectivity (0.452 ± 0.005 SEM), exceeding all other regions (all pairwise comparisons *P <* 0.001; largest effect: SSp vs. MO, Cohen’s *d* = 0.580; **Fig. 3F**). The remaining regions were comparatively similar (VIS: −0.034 ± 0.003 SEM; RSP: −0.135 ± 0.003 SEM; MO: −0.178 ± 0.004 SEM; all pairwise |*d*| *<* 0.17). This regional profile contrasted with the spatial tuning metrics: SSp neurons, despite exhibiting the weakest spatial consistency and lowest sparsity, showed the strongest distance-dependent functional coupling, while neurons in RSP and VIS exhibited high spatial consistency and sparsity but low distance-dependent functional coupling. Together, these analyses revealed a posterior-to-anterior gradient in spatial consistency and sparsity across dorsal cortex, alongside a dissociation between the spatial information content of individual neurons and the strength of local functional coupling.

### The neural manifold reflects the geometry of the physical space

To examine how population-level neural activity relates to spatial navigation, we performed principal component analysis (PCA) on the LSTFs. Because some cells could not always be tracked across all sessions, we implemented probabilistic PCA with missing data using an Expectation-Maximization approach [37, 31] (Methods A.8).

Performing PCA on the LSTFs across all sessions for each mouse, we found that the first three principal components traced out a saddle-shaped manifold that directly corresponded to the geometry of the figure-8 maze environment. Notably, this manifold was consistent across all mice (**Fig. 3B**), and there was also a clear differentiation in the activity at the central intersection of the figure-8 maze depending on whether mice approached from the left or right side, suggesting that cortical representations encoded both spatial location and directional context. The neural manifold also appeared to bend inwards on itself, bringing the two center locations and the two reward zones closer together in latent space, mirroring the left-right symmetry of the figure-8 maze.

These results demonstrated that the collective activity of neurons across multiple cortical regions formed a low-dimensional neural manifold whose geometry directly mirrored the geometry of the physical space the mice were navigating.

### Population dynamics systematically decorrelate over time

We next investigated whether population-level spatial representations remain stable or undergo systematic changes over the course of the 47-day recording period. To quantify representational stability, we computed cross-correlation matrices (CCMs) by measuring population activity similarity (Pearson correlation) at each spatial bin between all pairs of recording sessions across all mice (Methods A.12). Representational similarity was high at adjacent sessions (*r* = 0.466 ± 0.106 SD across mice) and declined substantially at maximum session separation (*r* = 0.117 ± 0.061 SD; **Fig. 4A-C**), confirming systematic decorrelation across the full recording period.

**Figure 4:**
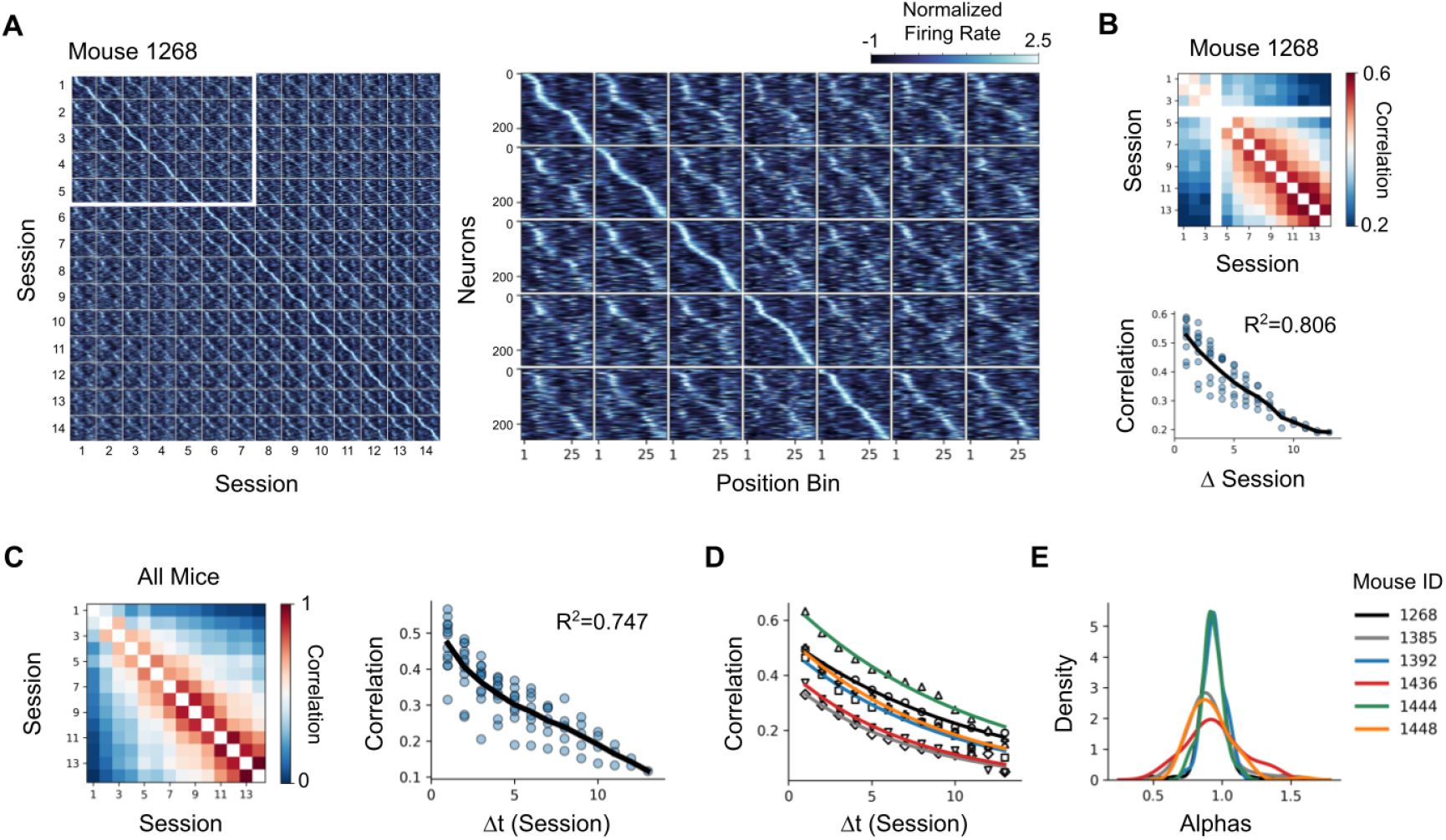
Population representations decorrelate across sessions. **(A)** Left: session-by-session sorted tuning matrix for mouse 1268, with cells ordered by peak tuning location within each diagonal block. Right: magnified view of five representative row segments from the full matrix. **(B)** Top: pairwise population correlation matrix for mouse 1268. Bottom: mean population correlation as a function of session separation (Δ*t*), with individual points representing correlations between session pairs at each lag, and the mean-per-Δ*t* curve shown in black. **(C)** Population correlation across sessions averaged across all mice. **(D)** Exponential curves fitted to the mean-per-Δ*t* curve for each mouse. **(E)** The distribution of multiplicative changes between all consecutive sessions. Values below 1 indicate a decrease in correlation, values above 1 indicate an increase in correlation.

To characterize the timescale of this decay, we fit an exponential function *ρ*(Δ*t*) = *A* exp(−Δ*t/τ*) to the off-diagonal session-pair correlations as a function of session separation Δ*t* (**Fig. 4D**; Methods A.13). Mice were imaged on a near-uniform schedule (mean inter-session gap ± SD: 3.5 ± 0.9 days, range 2–5 days; Methods A.3). The single-exponential model provided the best fit over linear (ΔAIC = 9.14 ± 3.35 SEM) and power-law (ΔAIC = 23.43 ± 2.00 SEM) alternatives, yielding a baseline similarity of *A* = 0.505 ± 0.042 SEM and a decorrelation timescale of *τ* = 9.51 ± 0.71 SEM sessions (*R*^2^ = 0.666 ± 0.042 SEM on individual session pairs; *R*^2^ = 0.978 ± 0.003 SEM on the mean-per-Δ*t* curve; all summary statistics across *n* = 6 mice).

An extended model incorporating an asymptotic offset (*ρ*(Δ*t*) = *A* exp(−Δ*t/τ*) + *c*) provided no improvement in fit (ΔAIC = 0.007 ± 0.943 SEM) with the fitted offset being small (*c* = +0.028 ± 0.016 SEM). Extrapolation suggested that population representations would con-tinue to decorrelate toward chance levels, with no persistent stable component. The distribution of session-to-session multiplicative changes (consecutive ratios of off-diagonal correlations along the same row of each CCM, **Fig. 4E**) was significantly below 1.0 in every mouse (one-sample *t*-test against *µ*_0_ = 1, one-sided, all *P <* 0.05), consistent with continuous, gradual decorrelation. Early sessions deviated from the exponential trend, reflecting initial learning dynamics that subsequently stabilized (Supplementary B.1 and B.2).

### Representational drift is not explained by behavioral change

The observed decorrelation could reflect drift in neural representations or changes in the animal’s behavior across sessions. With the mouse tethered to the robotically actuated head-stage, we were able to record the position, velocity, acceleration, and forces applied by the mice on the headstage. To evaluate the possible contribution of behavioral changes to changes in the neural representation, we constructed behavioral LSTFs alongside the neural LSTFs (**Fig. 5A**). Whereas neural LSTFs captured how population activity at each maze position is distributed across neurons, behavioral LSTFs captured how the animal’s observable behavior at each position is distributed across behavioral variables (Methods A.15). We then computed per-bin session-by-session CCMs from the behavioral LSTFs using the same procedure as for neural CCMs (**Fig. 5B,C**). This allowed us to compare how neural and behavioral representations changed over time at each position bin.

**Figure 5:**
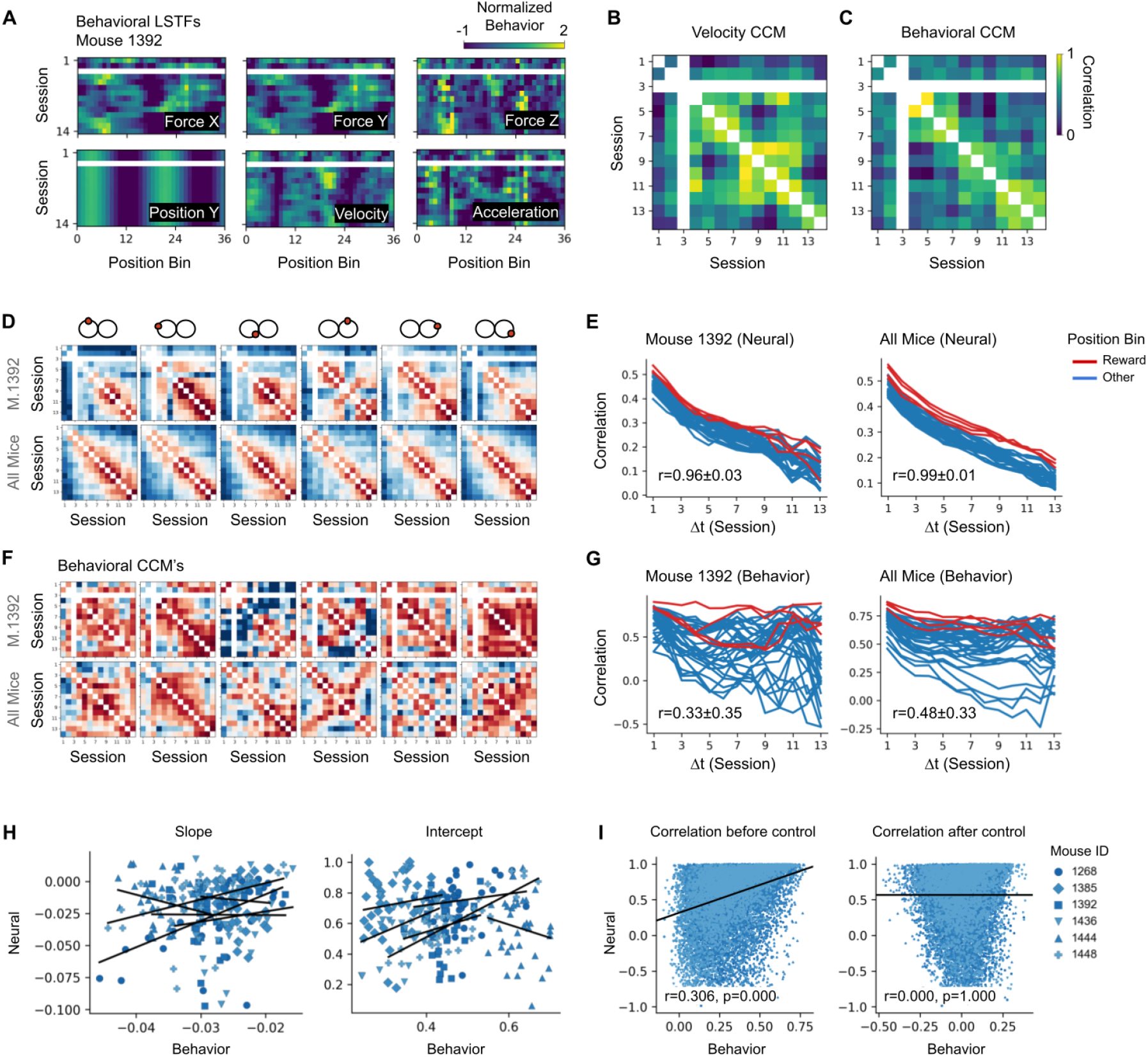
Representational drift and behavioral change. **(A)** Behavioral LSTFs computed in an identical manner to the neural LSTFs, shown for mouse 1392. **(B)** Session-by-session behavioral CCMs computed on a single behavior (velocity). **(C)** Session-by-session behavioral CCMs computed across all behavioral LSTFs for mouse 1392. **(D)** Neural CCMs at six representative position bins (4, 8, 15, 22, 26, 29 of 36; mouse 1392, top; across-mouse mean, bottom). **(E)** Per-bin neural drift curves (mouse 1392, left; across-mouse mean, right). **(F)** Behavioral CCMs at the same six position bins as in (D) (mouse 1392, top; across-mouse mean, bottom). **(G)** Per-bin behavioral drift curves (mouse 1392, left; across-mouse mean, right). **(H)** Comparison of neural and behavioral drift curves. Left: slope of behavioral drift versus slope of population correlation decay from (D-G). Right: intercept of behavioral drift versus intercept of population correlation decay from (D-G). **(I)** Correlation of behavioral and neural CCMs at each position bin before the behavioral control (left), and after the behavioral control (right). Black line shows linear regression fit.

Neural decorrelation was spatially uniform across position bins (**Fig. 5D,E**). If neural drift were driven by changes in behavior, we would expect drift to be strongest at the specific maze positions where behavior changed most. We therefore examined drift rates at individual position bins rather than averaged across the track. For each bin, we extracted the per-bin neural CCM entries as a function of Δ*t* and fit a linear regression of correlation against Δ*t*. The resulting per-bin slopes and intercepts were similar across all 36 positions (slope: −0.027 ± 0.007 SD across bins; intercept: 0.449 ± 0.115 SD; coefficient of variation (CV) for slope: 0.26; **Fig. 5E**). Behavioral decorrelation, by contrast, varied across the maze (**Fig. 5F,G**). Behavioral slopes were shallower on average and far more variable across positions (slope: −0.015 ± 0.027 SD; CV: 1.81; **Fig. 5G**), indicating that behavioral change was not uniform and instead was dependent on specific position bins. Correlations between neural and behavioral regression parameters (slope and intercept) were weak and varied in sign across mice (**Fig. 5H**; slope: *r* range −0.49 to 0.59; intercept: *r* range −0.16 to 0.80), indicating that the maze positions where behavior changed most were not the positions where neural representations drifted most.

To more directly control for behavioral contributions to neural decorrelation, we regressed the behavioral CCMs out of the neural CCMs at each spatial bin and recomputed the drift curves (Methods A.16). Neural decorrelation persisted after regressing out the behavioral contributions (*τ* = 11.35 ± 0.98 SEM sessions, *R*^2^ = 0.692 ± 0.042 SEM on individual session pairs; *R*^2^ = 0.969 ± 0.006 SEM on the mean-per-Δ*t* curve; compared with *τ* = 9.51 ± 0.71 SEM sessions before regression), confirming that the temporal changes in population codes were not principally explained by changes in the animal’s behavior. Per-bin neural and behavioral CCMs were correlated before regression (pooled across mice and bins, *r* = 0.306; per-mouse mean *r* = 0.357 ± 0.043 SEM), with the regression removing this shared variance (**Fig. 5I**). The remaining drift was slightly slower but nevertheless persisted throughout the recording period.

These analyses established that the neural decorrelation reported in the preceding sections reflected genuine representational drift: it proceeded at a uniform rate across the maze, persisted after regressing out behavioral variables, and was distinct from the position-specific changes observed in behavior.

### Representational drift is coordinated across cortical brain regions

We next asked whether different cortical regions follow distinct temporal dynamics reflecting their specialized functions, or instead share a common drift structure despite differences in baseline coding properties. We computed separate session-by-session CCMs for each region (RSP, VIS, MO, SSp) in each mouse (**Fig. 6A,B,C**). Adjacent-session correlations differed across regions in a manner consistent with the regional differences in spatial tuning described above (**Fig. 6C**): RSP showed the highest correlations (*r* = 0.501 ± 0.141 SD across mice), followed by VIS (0.486 ± 0.080), SSp (0.438 ± 0.077), and MO (0.404 ± 0.092). Despite these baseline differences, all four regions decorrelated on a similar timescale (mean ± SEM across mice): RSP (*τ* = 9.3 ± 0.6 sessions), VIS (9.1 ± 1.0), SSp (9.9 ± 1.1), and MO (9.5 ± 1.1). The CCMs themselves were also highly correlated across regions (mean *r* = 0.823 ± 0.128 SD across region pairs; **Fig. 6B**), indicating that the temporal profile of representational change was broadly similar across cortical areas.

**Figure 6:**
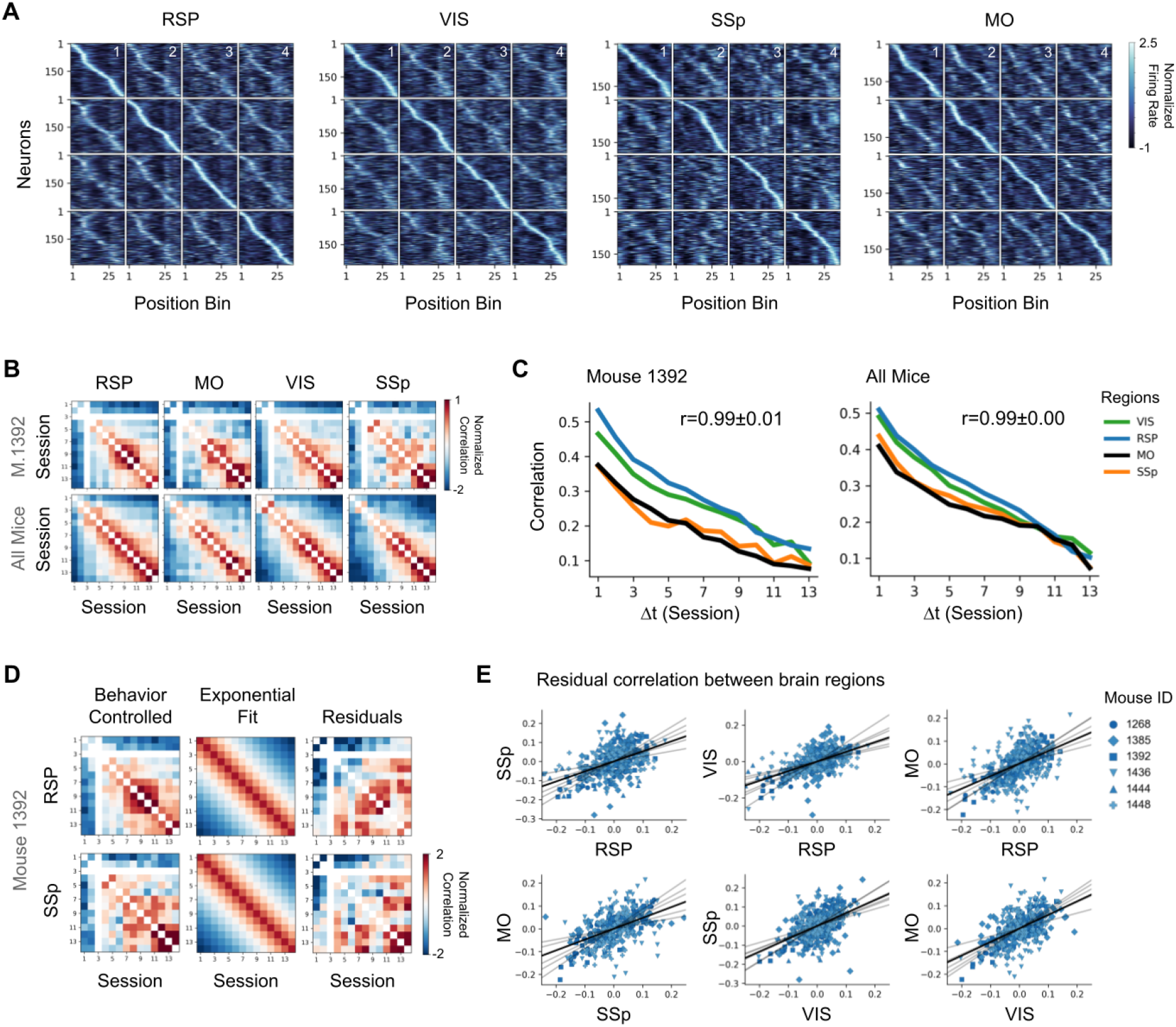
Representational drift across cortical regions. **(A)** Region-specific sorted tuning matrices for retrosplenial (RSP), visual (VIS), primary somatosensory (SSp), and motor (MO) cortices in mouse 1268. **(B)** Session-by-session population correlation matrices across cortical regions for mouse 1392 (top) and averaged across all mice (bottom). **(C)** Drift curves computed from the CCMs in (B) computed for each cortical region for mouse 1392 (left) and averaged across all mice (right). **(D)** Left: Behavior-controlled neural CCMs. Middle: the exponential fit to the neural CCMs in the left image. Right: the residuals computed by subtracting the exponential fit CCMs from the behavior-controlled neural CCMs. Shown in mouse 1392 across the RSP (top) and SSp (bottom). All CCMs are z-score normalized. **(E)** Correlation of the residuals from (D) across all pairwise regions, with all mice overlaid in each panel.

To test whether this similarity reflected coordinated drift or independent processes operating at comparable rates, we examined whether the residuals from each region’s expected decay were correlated across regions. We fit a single-exponential to each region’s drift curve and computed residuals (observed minus predicted correlation) for each session pair. These residuals capture how much each region drifted relative to its own expected trajectory at each session interval, after removing each region’s overall exponential decay. Correlated residuals would indicate that drift is coordinated session by session, while uncorrelated residuals would indicate that similar timescales arise from independent processes. We then correlated these residuals across all pairs of regions within each mouse (**Fig. 6D,E**; Methods A.14).

Cross-region residual correlations were positive in all 33 valid mouse-by-region-pair comparisons (3 of 36 excluded because one mouse lacked sufficient MO recordings), with a mean of *r* = 0.664 ± 0.029 SEM and consistent values across animals (per-mouse mean range 0.52 to 0.86; **Fig. 6E**). To account for the fact that each mouse contributed multiple region-pair comparisons, we fit a linear mixed-effects model on Fisher-*z*-transformed correlations with mouse as a random intercept, yielding a population-level *r* = 0.700 (95% CI [0.550, 0.806]; Methods A.14). All 33 observed correlations also exceeded a per-comparison shuffle null at *P* < 0.01, and a binomial sign test on the count of positives gave *P* = 1.2 × 10^*−*10^. Coordination did not depend on specific regional pairings (RSP–VIS: *r* = 0.714 ± 0.061; RSP–SSp: 0.613 ± 0.063; RSP–MO: 0.669 ± 0.093; VIS–SSp: 0.662 ± 0.074; VIS–MO: 0.679 ± 0.073; SSp–MO: 0.650 ± 0.087; mean ± SEM across mice).

This coordination persisted under three robustness controls. First, regressing the behavioral CCMs out of the neural CCMs preserved the coordination (mean *r* = 0.579 ± 0.032 SEM; 33/33 positive), ruling out shared behavioral fluctuations as the source. Second, excluding the first three sessions, which deviate from the exponential trend during early-task learning, preserved the coordination as well (mean *r* = 0.557 ± 0.040 SEM; 33/33 positive). Third, applying both the behavioral control and the early-session exclusion in combination still preserved the coordination (mean *r* = 0.487 ± 0.040 SEM; 33/33 positive), indicating that drift was coordinated throughout the recording period and not solely during behavioral familiarization.

### Population-level spatial representations are preserved through geometric invariance

The preceding analyses established that single-neuron spatial representations decorrelated over time across all cortical regions. Yet the PCA analysis revealed a shared manifold structure across sessions, raising the question of how population-level representations could remain functional despite ongoing changes in individual neurons.

To assess the stability of population-level spatial structure, we computed position-by-position cross-correlation matrices (CCMs) that capture the co-variation between spatial locations as encoded by the neural population. For each session, we calculated the Pearson correlation between population activity vectors at every pair of spatial bins (*N* = 36), yielding a 36 × 36 matrix representing the geometric similarity structure of the population code (**Fig. 7A,B**).

**Figure 7:**
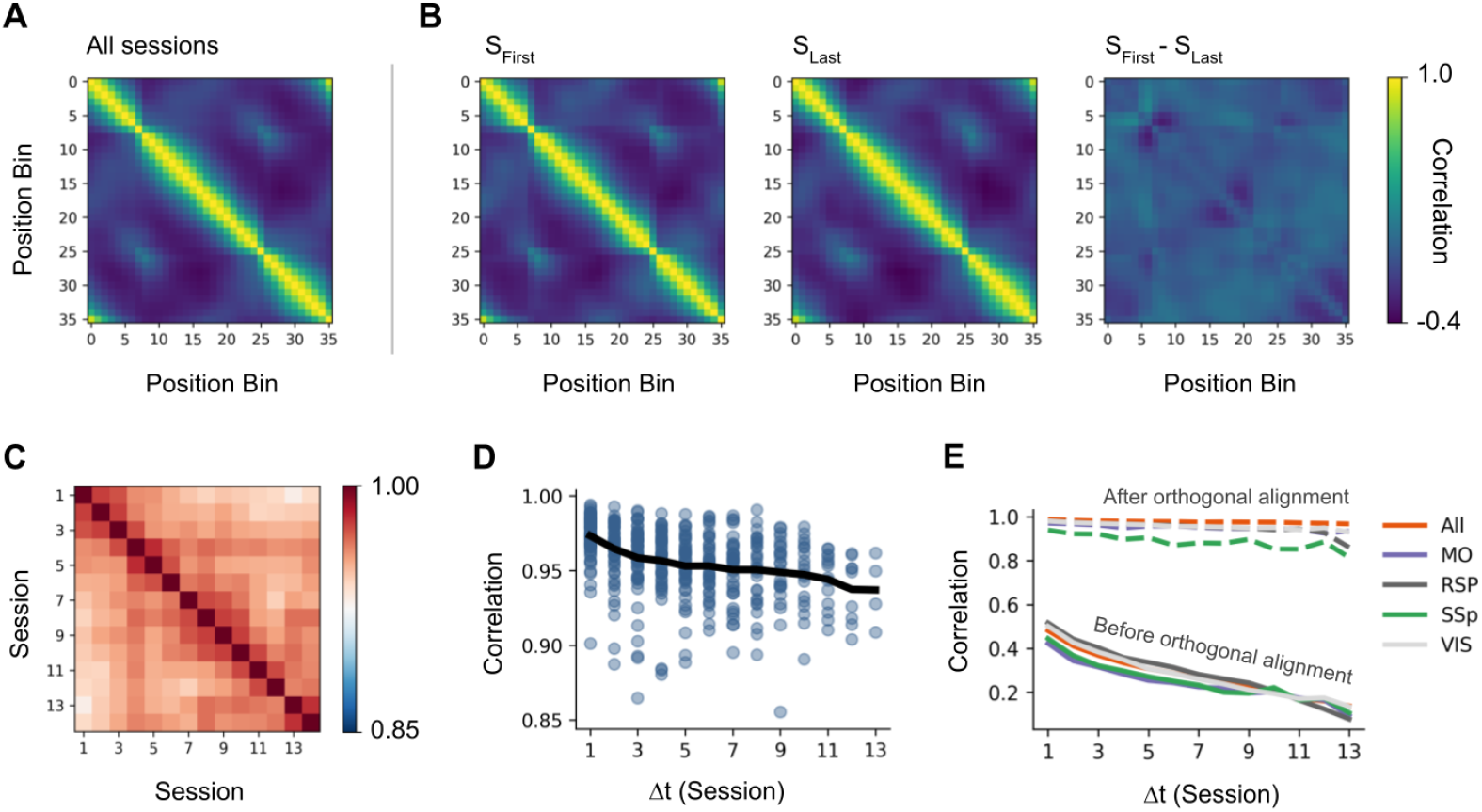
Preserved population geometry. **(A)** Position-bin-by-position-bin correlation matrix averaged across all sessions and mice. **(B)** Position-bin-by-position-bin correlation matrices for the first (left) and last (middle) sessions, averaged across mice, and their residual (right). **(C)** Cross-correlation matrix computed between the session-averaged position-bin-by-position-bin matrices across sessions. **(D)** Pairwise correlations derived from (C), with the mean curve shown in black. **(E)** Correlation between population activity vectors before and after orthogonal alignment, averaged across mice, shown for all regions combined (All) and for individual regions (RSP, VIS, MO, SSp).

These spatial co-variation matrices remained highly stable across sessions despite ongoing single-neuron drift. Pairwise comparisons between non-overlapping sessions yielded mean *r* = 0.957 ± 0.023 SD across pairs (*N* = 940 independent session pairs across all mice), far exceeding chance levels established by shuffled controls in which spatial bin labels were randomly permuted (mean *r* = 0.210 ± 0.048; Cohen’s *d* = 16.010). Correlation declined only modestly over time: mean *r* = 0.973 ± 0.015 for adjacent sessions decreased to *r* = 0.939 ± 0.021 at maximum session separation, representing a 3.5% decline over the full 47-day recording period (**Fig. 7C,D**).

Preserved co-variation structure under changing neural responses implied that the transformation between sessions was approximately orthogonal, provided that population norms scaled approximately uniformly across sessions (Supplementary B.3 for a formal derivation). To test this, we aligned population activity between all pairs of sessions using orthogonal Procrustes analysis, which finds the optimal rotation (and reflection) mapping one session’s population vectors onto another’s (Methods A.17). Before alignment, the correlation between population activity vectors was moderate (*r* = 0.334 ± 0.135 SD across pairs), reflecting the cumulative effect of single-neuron drift. After orthogonal alignment, correlations increased to *r* = 0.979 ± 0.009 with low normalized residual error (*ϵ* = 0.258 ± 0.080), both substantially exceeding shuffled controls (shuffle: *r* = 0.497 ± 0.051, *ϵ* = 1.035 ± 0.098). The results before and after orthogonal alignment are shown in **Fig. 7E**. Thus, while individual neurons continuously changed which spatial locations they encoded, the relative relationships between locations in the population code remained stable, with representational change between sessions well described by rotation-like transformations that preserved the geometric structure of the spatial code.

## Discussion

Using longitudinal widefield calcium imaging enabled by a cranial exoskeleton, we tracked over 110,000 unique layer 2/3 neurons across four cortical regions spanning 47 days as mice navigated a familiar figure-8 environment. Despite regional differences in spatial coding density, population representations across retrosplenial, visual, somatosensory, and motor cortices decorrelated over time with similar exponential timescales. Critically, session-specific fluctuations in representational similarity were correlated across all region pairs, suggesting that drift may not be an independent process within each area but may instead be coordinated across the dorsal cortex. Despite this coordinated drift, population-level geometric structure was preserved: the relative relationships between spatial representations remained stable across sessions, with inter-session transformations well described by an orthogonal transformation of the population code.

Spatial coding properties varied systematically across the dorsal cortex, following a posterior-to-anterior gradient in both spatial consistency and sparsity. RSP and VIS contained the highest proportions of spatially tuned neurons, consistent with established roles for these regions in integrating visual landmarks with internal spatial representations during navigation [39, 25, 1]. This gradient was not mirrored in local functional connectivity: primary somatosensory cortex, despite exhibiting the lowest spatial consistency and sparsity, showed the strongest distance-dependent functional coupling among neurons within a hemisphere. This dissociation would indicate that spatial coding and local functional coupling reflect distinct organizational features across dorsal cortex, with SSp exhibiting strong distance-dependent co-firing among neighboring neurons despite contributing least to spatial coding of the maze environment. Conversely, neurons in posterior regions such as RSP and VIS showed the highest spatial consistency and sparsity but the weakest local functional coupling.

RSP engagement varied across animals: four of six mice displayed high RSP spatial consistency throughout the recording period, while the remaining two showed less reliable RSP spatial tuning. The constrained movement imposed by the cranial exoskeleton, which guided mice along a fixed trajectory, may have permitted multiple navigational solutions: mice could have anchored their position estimates using visual landmark information routed through retrosple-nial circuitry, or alternatively solved the task through procedural strategies relying on learned motor and proprioceptive cues that do not require posterior spatial integration [39, 14, 29].

Having characterized the spatial tuning properties of individual neurons across the dorsal cortex, we next asked how these representations behaved at the population level across the 47-day recording period. All four cortical areas showed exponential decorrelation with overlapping timescales (*τ* range across regions: 9.1–9.9 sessions, mean across mice), and session-to-session deviations from each region’s expected decay trajectory were positively correlated across all pairwise region combinations. This coordination persisted after regressing out behavioral variables measured simultaneously via the exoskeleton (position, velocity, acceleration, and applied force), and after excluding early sessions to control for task-acquisition learning effects, controlling for shared behavioral fluctuations and early-session learning. These findings were inconsistent with drift emerging fully independently within each cortical area, and instead suggested that the temporal dynamics of representational change may be coupled across the dorsal cortex.

Despite continuous drift in single-neuron tuning, population-level geometric structure was highly stable over the 47-day recording period. Position-by-position covariance matrices, which captured the relative similarity between spatial locations as encoded by the neural population, were correlated across sessions at *r* = 0.957, declining by only 3.5% from adjacent to maximum session separation. Orthogonal Procrustes alignment showed that the transformation between sessions’ population codes was well described by rotation and reflection in neural state space, increasing cross-session population correlation from *r* = 0.334 before alignment to *r* = 0.979 after alignment. This was not merely a descriptive observation: as we formally derive (Supplementary B.3), preserved covariance structure under a linear inter-session transformation implied the transformation was approximately orthogonal, given uniform norm scaling of population vectors. Such transformations preserved the angles and distances between population embeddings, maintaining the functional geometry of the spatial code even as individual neurons changed their tuning. This provided empirical grounding for theoretical proposals that geometric invariance of population codes may support stable computation despite continuous single-neuron drift, without requiring permanence of individual tuning properties [11, 33, 21, 5].

## Methods

### A.1 Surgery

All animal experiments were approved by the University of Minnesota’s Institutional Animal Care and Use Committee (IACUC). The cranial implants for the robot-rodent interface were 3D printed from photo-curing polymer and designed to conform specifically to the dorsal surface of the mouse’s skull. The implant structure included three 0-80 tapped holes for chronic headpost fixation and two additional holes over the occipital plate to secure the device using skull screws. To facilitate mesoscale imaging, the implant featured a 6 × 6 mm glass plug that compressed the dorsal cortex into a single flat imaging plane. This plug was constructed from a 0.5 mm thick fused quartz top glass and a 1 mm thick bottom glass slide, which were bonded to each other and the implant using UV-curable optical glue. The surgical procedure began with the administration of anesthesia using 1–5% isoflurane, followed by buprenorphine and meloxicam for analgesia and inflammation control. After securing the mouse in a stereotax, the scalp was excised and craniotomies were performed using a high-speed dental drill. The sterilized implant was then placed on the skull, secured with screws, and adhered using dental cement. Once the cement was cured, the headpost was attached with screws, and a plastic lid was placed to protect the glass surface. Mice were monitored on a heated pad until they were fully ambulatory and were given a recovery period of at least 14 days before behavioral acclimation began.

### A.2 Exoskeleton operation

The cranial exoskeleton was a three-armed delta robot designed to provide translational motion in the x, y, and z axes, integrated with a motorized goniometer that enabled rotational control over pitch, roll, and yaw [17]. Mice were coupled to the headstage via a kinematic clamping mechanism attached to a chronically implanted titanium headpost. Using a six-axis force sensor, the system measured the forces applied by the mouse to drive an admittance control law, allowing the animal to control its own velocity and acceleration within the limits of the system. This robotic platform supported a high-performance mesoscale imaging payload, specifically a custom-built macroscope designed to record Ca^2+^ activities from thousands of neuronal sources across a wide 8.6 × 6.6 mm field of view.

The experimental protocol began with a multi-stage habituation process, where mice were first head-fixed on a cylindrical treadmill until they could locomote comfortably for at least 10 minutes. Following this initial acclimation, mice were habituated to navigating a figure-8 maze while learning to maneuver the exoskeleton. During the actual experiment, the robot was positioned over a cylindrical treadmill, and the mouse was docked to the headstage via a kinematic clamping mechanism. The researcher then adjusted the headstage optics to account for mouse-to-mouse variability while the mouse paced on the treadmill to ensure a stable interface before imaging began. Once the system was configured, the mouse was ushered into the figure-8 maze environment to begin the behavioral session. Within the maze, the mouse actively controlled its velocity and acceleration by applying cranial forces that the exoskeleton’s admittance control law translated into movement, and a trajectory control algorithm ensured the mouse stayed on the defined trajectory of the figure-8 maze. To ensure precise alignment between imaging and behavioral data, a TTL signal synchronized the LED driver with the exoskeleton’s telemetry, aligning the image series (saved in TIFF format) with the robot’s positional data. The session continued until the mouse ceased to perform or reached the specific trial criteria, at which point the subject was removed from the headstage and returned to its home cage.

### A.3 Mouse selection and habituation

For selection, mice were first habituated to running on a wheel for 3 sessions, with each session ranging from 1–7 minutes in length. Imaging was then done on the mice while they ran on the wheel using the cranial exoskeleton, after which 6 mice were selected from a cohort of 15 based on the quality of GCaMP6s calcium indicator expression.

Before running the mice through the figure-8 maze environment, they were once again acclimated to the maze without the cranial exoskeleton during 4 habituation sessions that each lasted 6 minutes in length. Once the mice were habituated, they were attached to the cranial exoskeleton 3–4 days later to start the experiment. All mice were imaged on the same schedule of 14 sessions on days 1, 5, 7, 9, 12, 15, 19, 23, 26, 30, 33, 37, 42, and 47, with one exception: mouse 1268 was imaged one day earlier than the other mice on session 8 (day 22 instead of day 23). Inter-session gaps were therefore tightly distributed (mean ± SD: 3.5 ± 0.9 days, range 2–5 days). Three sessions of data were lost to recording failures (mouse 1392 session 3, mouse 1436 session 5, mouse 1385 session 13) and were excluded from analysis.

### A.4 Calcium imaging with the cranial exoskeleton

Imaging was performed using a custom-built macroscope integrated within the cranial exoskeleton developed previously [17]. The macroscope consists of an excitation light system and monochrome camera system housed in adjustable mounting brackets allowing proper alignment with the implant. The camera system can image an 8.6 × 6.6 mm field of view at a design resolution of 3.45 × 3.45 µm (pixel size). A cranial implant, adapted from Hattori and Komiyama [16], was surgically implanted over the mouse’s brain to optically expose 35 mm^2^ of the dorsal cortex. Imaging was performed on double transgenic mice [8] (Cux2Cre-ERT2 [10] × Ai163 [6]) sparsely expressing Ca^2+^ indicator GCaMP6s in layer 2/3 excitatory neurons [18]. Data acquisition was performed on a PC using software from the camera supplier that saved an image series in TIFF format. A synchronization signal linked the image series data and exoskeleton data for future analysis.

### A.5 Calcium imaging processing

Raw imaging data was processed using Suite2p (v0.11.1) [26] at an acquisition rate of 10 Hz. Motion correction was performed using non-rigid registration with a block size of 128 × 128 pixels, a maximum shift of 10% of the field of view, and a smoothing sigma of 4.0 pixels. Neurons were detected using sparse mode with a cell diameter of 12 pixels, a threshold scaling factor of 1.0, and a maximum allowed ROI overlap of 75%. Raw fluorescence (*F*) and neuropil (*F*_neu_) traces were extracted for each detected cell with an inner neuropil radius of 2 pixels and a minimum neuropil pixel count of 350. ΔF/F was computed by first estimating a slow baseline of *F*_neu_ using a sliding window percentile filter (window = 800 frames, 33rd percentile), then computing the neuropil-corrected signal as (*F* − *F*_neu_) / mean(*F*_neu_ baseline). A second baseline, estimated by applying the same filter to the resulting ΔF/F, was then subtracted to remove residual low-frequency drift. Cells were then manually validated and filtered based on the spatiotemporal properties of their extracted footprints and ΔF/F traces.

### A.6 Cell registration

Our cell registration algorithm consisted of three sequential stages: (i) cross-session image alignment, (ii) pairwise similarity scoring, and (iii) iterative cluster assignment, each applied independently per mouse. For each session, mean fluorescence projection images and a composite image of all cell footprints were constructed from Suite2p output (**Fig. 2C**). Mean projections were preprocessed to enhance anatomical contrast, and cell footprints were super-imposed to produce a single image used for registration. All sessions were registered to the temporally middle session for each mouse. Registration proceeded in two stages: an initial projective transformation was estimated from matched SURF features, followed by a refinement step using matched cell centroid pairs from the initial alignment. The final composite transformation was applied to register each session’s cell footprints and centroids into the reference frame.

For each pair of sessions, candidate cell matches were identified by searching within a 34.5 µm radius of each registered cell centroid, and spatial correlation between cell footprints was computed as their normalized inner product. Adapting the framework of Sheintuch et al. [36], a per-mouse probabilistic model was fit to estimate the probability *P*_same_ that a candidate pair represents the same neuron as a function of spatial correlation. Nearest-neighbor and other-neighbor correlation distributions were modeled parametrically: same-cell correlations with a lognormal distribution and different-cell correlations with a beta distribution, weighted by a prior same-cell probability *W*_same_ estimated from the marginal centroid distance distribution. This yielded a 1D *P*_same_ score as a function of spatial correlation, fit independently for each mouse.

Cell identities across sessions were resolved by greedy clustering seeded from the earliest session, followed by iterative global refinement. In the greedy step, for each new session the globally maximum *P*_same_ pair was matched and removed from consideration iteratively until no remaining pair exceeded *P*_same_ ≥ 0.5. Unmatched cells were initialized as new clusters. Clusters were then iteratively refined by reassigning cells to whichever cluster yielded the highest *P*_same_, resolving conflicts by displacing the lower-scoring occupant, until convergence. Clusters appearing in only a single session were discarded. Following cross-session registration, each mouse’s reference frame was aligned to the Allen Mouse Brain Atlas, enabling each tracked cell to be assigned to a cortical region (RSP, VIS, SSp, or MO) for the region-specific analyses described below.

### A.7 Lapwise spatial tuning functions

Lapwise spatial tuning functions (LSTFs) were computed from the ΔF/F traces of each recorded neuron across all sessions. Raw ΔF/F traces were first smoothed using a 1D Gaussian filter (*σ* = 0.75 seconds, equivalent to 7.5 samples at 10 Hz acquisition). The linearized position of the mouse was discretized into *N* = 36 equally sized spatial bins (∼4.45 cm each), spanning the full length of the figure-8 maze track. Timepoints at which the mouse velocity fell below 0.39 cm/s were excluded to restrict the analysis to active locomotion epochs (threshold selected by manual inspection of the velocity distribution to separate stationary from moving epochs).

For each lap, the mean ΔF/F of each neuron was computed within each occupied spatial bin, yielding a per-lap tuning function of shape (bins × neurons). LSTFs were organized as a three-dimensional tensor of shape (laps × bins × neurons). Tuning functions were subsequently z-score normalized within each session across neurons to account for session-to-session variability in baseline fluorescence.

### A.8 Probabilistic PCA

To visualize population dynamics while handling incomplete cell tracking across sessions, we implemented probabilistic PCA with missing data [37, 31]. Since individual neurons could not always be tracked across all recording sessions, the spatial tuning function matrix contained NaN values for untrackable cells in specific sessions. The probabilistic framework treats missing data as latent variables that can be marginalized out, enabling dimensionality reduction from incomplete observations.

We retained *k* = 4 latent components per mouse. Loadings *W* and the mean *µ* were estimated by standard PCA on the across-lap mean spatial tuning functions, after z-score normalization and spatial smoothing (*σ* = 1.0 bins). Averaging across laps reduced missing values in the training set to a negligible fraction. With *W* and *µ* fixed, individual-lap projections were then obtained by inferring the latent score *z* for each lap from its observed entries only.

We treat each sample *x* as drawn from the standard probabilistic PCA model:

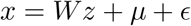

where *W* ∈ ℝ^*n×k*^ contains the principal components, *z* ∈ ℝ^*k*^ are the latent scores, *µ* is the mean, and *ϵ* is isotropic Gaussian noise. For samples with missing dimensions, we estimated the latent representation *z* from the observed dimensions only by solving:

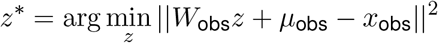

where the subscript “obs” denotes the subset of observed (non-NaN) dimensions, with closed-form solution

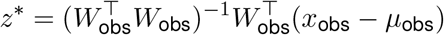

This corresponds to the zero-noise limit of the probabilistic PCA E-step [37] and was sufficient for the visualizations in this paper, where *W* is set from the dense training fit and not iteratively re-estimated.

For cross-mouse visualization (Fig. 3B), each mouse’s 4-component PC trajectory was aligned to a reference mouse by fitting and applying a linear map between corresponding trajectory entries.

### A.9 Spatial consistency score

Spatial consistency was quantified for each neuron using a leave-one-out cross-validation scheme across laps. For each fold, the mean tuning function was computed from all training laps and correlated with the held-out test lap using Pearson correlation. The consistency score was defined as the average correlation coefficient across all folds. Laps containing NaN values were excluded from analysis, and correlation coefficients that were NaN were set to zero. This metric quantifies the trial-to-trial stability of each neuron’s spatial tuning pattern.

Mathematically, for neuron *j* with tuning functions *f*_1_, *f*_2_, …, *f*_*n*_ across *n* laps, the spatial consistency *C*_*j*_ is defined as

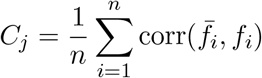

where 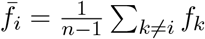 is the mean tuning function excluding lap *i*, and corr(·, ·) is the Pearson correlation.

### A.10 Spatial sparsity index

To quantify the spatial sparsity of individual neurons, we computed a sparsity index from each cell’s spatial tuning function. This metric captures how sparsely a neuron responds across spatial locations, with higher values indicating sharper spatial tuning.

For each neuron, spatial tuning functions were first normalized to form a probability distribution by subtracting the minimum response and dividing by the sum across all spatial bins. The sparsity index *S* was then computed as:

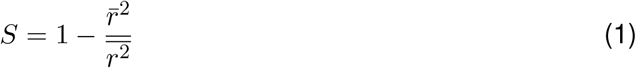

where 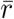 denotes the mean response across spatial bins and 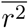 denotes the mean squared response. This formulation corresponds to a modified lifetime sparseness metric [30, 40], which ranges from 0 (uniform response across all locations) to 1 − 1*/N* (response confined to a single location). Neurons with tuning functions containing invalid values or zero variance were excluded from analysis.

For neurons tracked across multiple sessions, we computed the sparsity index independently for each session and reported the mean score across sessions. The within-mouse z-scoring and mixed-effects model used for region comparisons are described in Methods A.18.

### A.11 Functional connectivity

To examine how functional connectivity relates to anatomical organization across the dorsal cortex, we quantified the relationship between pairwise neural correlations and the physical distance between cells. This metric captures the intuitive notion that neurons with strong local connectivity should show higher correlations with nearby cells compared to distant cells. For each neuron *i*, we computed Pearson correlations *ρ*_*ij*_ from raw ΔF/F traces with all other simultaneously recorded neurons *j* within the same hemisphere, along with the corresponding Euclidean distances *d*_*ij*_ between cell centroids. The distance-dependent connectivity score for neuron *i* was then defined as:

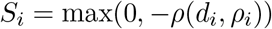

where *d*_*i*_ = {*d*_*ij*_: *j* ≠ *i*} and *ρ*_*i*_ = {*ρ*_*ij*_: *j* ≠ *i*} represent the vectors of distances and correlations for neuron *i*, respectively. The negative sign ensures that neurons exhibiting stronger local connectivity (negative correlation between distance and functional connectivity) receive higher scores, while the maximum operation with zero clips cases where distant neurons are more correlated than nearby neurons.

This analysis was performed on the ΔF/F traces for each session independently, without requiring cell tracking across sessions, resulting in a larger total cell count (*n* = 312,648 cells from 6 mice) that includes some cells recorded multiple times across sessions. For region comparisons, functional connectivity scores were z-score normalized within each mouse before pooling, following the convention described in Methods A.18.

### A.12 Cross-session correlation analysis

To quantify changes in neural population dynamics across sessions, we computed Pearson correlations between session-averaged LSTFs across all pairwise sessions. Specifically, letting 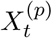 denote the mean population code at position bin *p* ∈ [1, 36] and session *t*, we defined the correlation between sessions *i* and *j* at position *p* as:

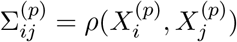

where *ρ*(*x, y*) is the Pearson correlation between *x* and *y*. Averaging across positions yields a session-wide measure of similarity:

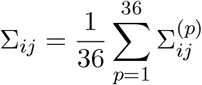

From Σ_*ij*_, we constructed decorrelation curves by plotting representational similarity as a function of session separation Δ*t* = |*i* − *j*|.

### A.13 Exponential decay fitting

Decorrelation curves were modeled as a single exponential decay, *ρ*(Δ*t*) = *A* exp(−Δ*t/τ*), where Δ*t* = |*i* − *j*| is the session separation, *A* is the baseline correlation, and *τ* is the decorrelation timescale (in sessions). Fits were performed independently for each mouse and each cortical region.

For each (mouse, region), we extracted all valid off-diagonal entries of the session-by-session correlation matrix Σ (Methods A.12), excluding pairs with NaN correlations (e.g., from sessions with insufficient cell tracking). Parameters (*A, τ*) were estimated by non-linear least squares (`scipy.optimize.least_squares`), with *τ* parameterized as log(*τ*) to enforce positivity and initial values *A*_0_ = max_(*i,j*)_ *ρ*_*ij*_ and 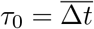.

For the cross-region coordination analysis (Methods A.14), residuals were computed as Σ_*ij*_ − *A* exp(−|*i* − *j*|*/τ*) at each observed session pair.

Goodness-of-fit was reported as *R*^2^ on individual session pairs and on the mean-per-Δ*t* curve (averaging correlations within each unique Δ*t* before computing *R*^2^). The single-exponential model was compared against linear (*ρ* = *a*Δ*t* + *b*), power-law (*ρ* = *A*Δ*t*^*−β*^), and offset-exponential (*ρ* = *A* exp(−Δ*t/τ*) + *c*) alternatives using the Akaike Information Criterion on each mouse’s mean-curve fit.

### A.14 Cross-region coordination test

To test whether session-pair-specific deviations from the expected exponential decay were shared across cortical regions, we computed Pearson correlations between regional residual matrices within each mouse. For each pair of regions in each mouse, we extracted the upper-triangular off-diagonal entries of the residuals (Methods A.13), retained only entries that were non-missing in both regions, and computed the Pearson correlation between these vectors. With four regions and six mice, this yielded up to 6 region-pairs × 6 mice = 36 mouse × region-pair correlations, of which 33 were valid (mouse 1444 lacked sufficient recordings in motor cortex, which removed 3 pairs).

We assessed coordination using two complementary tests. First, we constructed a permutation null for each comparison by shuffling the session-pair indices of one residual vector before correlating, repeated *B* = 10,000 times per comparison. The empirical *P* -value was computed as *P* = *k/B*, where *k* is the count of shuffles with |*r*_shuffle_|≥|*r*_obs_|, giving a precision floor of 1*/B* = 10^*−*4^. This procedure preserved the marginal distribution of each residual vector while breaking the cross-region session-pair alignment. Because the test fixes the marginal residual distribution per region, it isolates alignment from chance but cannot exclude shared session-level signals (e.g., recording-quality fluctuations, global cortical state) that produce genuine cross-region alignment.

Second, to account for clustering of comparisons within mice, we fit a linear mixed-effects model on Fisher-*z* transformed residual correlations (*z* = tanh^*−*1^(*r*), with *r* clipped to ±0.999 for numerical stability) with mouse as a random intercept. The intercept estimates the population-level Fisher-*z* correlation. The corresponding Pearson correlation and 95% CI were back-transformed from the Wald interval on the intercept. Models were fit by restricted maximum likelihood (REML) via `statsmodels.formula.api.mixedlm` in Python with *n*_obs_ = 33 region-pair correlations clustered in *n*_clusters_ = 6 mice.

As a non-parametric supplementary check, we report the count of positive correlations with an associated one-sided exact binomial sign test (*H*_0_: *P* (*r >* 0) = 0.5).

We additionally evaluated coordination under three robustness controls, each applied before computing the regional residuals: regressing the behavioral CCMs out of the neural CCMs at each spatial bin (Methods A.16), excluding the first three canonical sessions from each mouse’s session-by-session matrix to remove early-task learning dynamics, and applying both controls in combination. In each case, the per-region exponential fits and per-pair Pearson residual correlations were recomputed using the same procedure described above.

### A.15 Behavioral lapwise spatial tuning functions

To construct behavioral analogs of the neural LSTFs, we recorded four distinct behavioral signals during each lap of navigation: mouse position (2D, *x* and *y* in the maze plane), velocity (1D magnitude), acceleration (1D magnitude), and the forces applied by the mouse on the headstage (3D, the *x, y, z* components from the six-axis force sensor). Concatenated, these yielded *K* = 7 behavioral channels per timepoint. For each lap, behavioral measurements were averaged within the same 36 spatial bins used for neural data, excluding periods of immobility (*<*0.39 cm/s). This yielded a behavioral tuning function for each lap. Behavioral tuning functions were z-score normalized within each session independently for each variable. The resulting behavioral LSTFs were organized as a three-dimensional tensor (laps × spatial bins × *K* behavioral channels), paralleling the structure of the neural LSTFs (laps × spatial × bins neurons). Furthermore, we applied the same level of Gaussian smoothing on the behavioral data as the neural data (*σ* = 0.75 seconds).

Behavioral CCMs were computed from the behavioral LSTFs using the same procedure as for neural CCMs, ensuring that neural and behavioral drift curves are computed in exactly the same manner, and that differences between neural and behavioral drift curves reflect differences in the underlying signals rather than methodological differences in how the CCMs were computed.

### A.16 Behavioral regression control

To control for the possibility that neural decorrelation was driven by changes in behavior, we regressed the behavioral CCMs out of the neural CCMs independently at each spatial bin. For each bin *p*, we fit a linear regression predicting 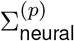 from 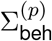 across all session pairs (*i, j*), yielding a slope 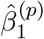 and intercept 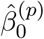. We then removed the behavioral contribution while preserving the baseline (mean) neural correlation level:

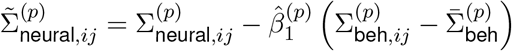

where 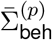 is the mean of the behavioral CCM across all session pairs at bin *p*. By subtracting the demeaned behavioral predictor scaled by the regression slope, this procedure removes variance in the neural CCM that is linearly predicted by behavioral change while preserving the overall scale of neural correlations. Session pairs with missing data in either CCM were excluded.

### A.17 Procrustes analysis

We performed orthogonal Procrustes analysis to determine whether orthogonal transformations could optimally align population activity patterns between sessions. For each pair of sessions, we computed neural activity matrices *X*_*t*_ and 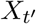 (spatial bins × neurons), where each row is the mean population vector at one of the 36 spatial bins. After mean-centering across bins, we found the orthogonal matrix *Q* ∈ ℝ^*n×n*^ (*n* = number of neurons tracked across both sessions) minimizing 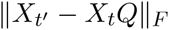, computed via the standard SVD solution. *Q* rotates (and possibly reflects) the population vector at each bin in neuron space, providing the orthogonal transformation framework derived in Supplementary B.3. We quantified alignment quality as the mean across spatial bins of the Pearson correlation between corresponding rows of 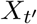 and *X*_*t*_*Q*, and as the normalized Frobenius residual 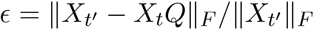.

### A.18 Statistical analysis

All statistical analyses were performed in Python 3.9 using `numpy` (1.24), `scipy` (1.13), and `statsmodels` (0.13). Throughout the paper, ± values denote either the standard deviation (SD) when summarizing variability across observations (e.g., across mice, position bins, or session pairs) or the standard error of the mean (SEM) when reporting the precision of fitted parameters or group-level means. Each ± value is labeled explicitly inline. For region comparisons of cell-level metrics (spatial consistency, sparsity, and functional connectivity), scores were z-score normalized within each mouse (subtract mouse mean, divide by mouse standard deviation) prior to pooling across mice, removing between-mouse differences in baseline. Regional differences were then assessed with a linear mixed-effects model fit by restricted maximum likelihood via `statsmodels.formula.api.mixedlm` with region as a fixed effect and mouse as a random intercept, and pairwise comparisons across the four cortical regions (6 pairs) were Tukey HSD-corrected. Permutation tests used *B* = 10,000 shuffles, with *P* computed as the proportion of shuffles meeting the test criterion (precision floor 1*/B* = 10^*−*4^). Model comparisons for decay-curve fits used the Akaike Information Criterion, with ΔAIC reported relative to the single-exponential model.

## Author contributions

R.P. designed and ran data analyses. M.F. developed the cell tracking algorithm. J.H. and T.B. ran mouse experiments. I.H. performed mouse surgeries. A.D.R. and S.B.K. performed supervision and provided feedback. S.B.K. provided funding acquisition. R.P. and S.B.K. wrote the manuscript.

## Acknowledgments

BRAIN Initiative grants RF1NS113287 and RF1NS126044. R.P. was supported by the Pecot Fellowship awarded by the McKnight Foundation, and MN Robotics Institute.

## Code & data availability

All code for cell extraction, data analysis, and reproducibility, along with the underlying data, will be made publicly available upon publication.

## B Supplementary

### B.1 Spatial consistency increases with time

One repeated pattern we observed in the single-cell LSTFs was that cell tuning would tighten and/or become more consistent over time (**Supplementary Fig. 3**). To quantify this change in spatial representations across the 47-day recording period, we correlated individual lap tuning functions with each neuron’s mean tuning across all sessions, and observed the change in correlation across time. The lap-wise correlation increased from early sessions (session 1: 0.324 ± 0.059 SD across mice) to late sessions (session 14: 0.435 ± 0.074) (**Supplementary Fig. 3B**).

This increase in consistency was observed across all cortical regions, and in each individual mouse. From session 1 to session 14, across all mice, visual cortex consistency increased from 0.369 ± 0.059 SD to 0.458 ± 0.064, retrosplenial cortex from 0.324 ± 0.085 to 0.452 ± 0.090, somatosensory cortex from 0.320 ± 0.058 to 0.440 ± 0.065, and motor cortex from 0.315 ± 0.066 to 0.414 ± 0.047. The consistent increase across all examined regions indicated that spatial tuning became more reliable with continued experience in this behavioral paradigm.

### B.2 Linear decoding of mouse position

To assess whether mouse position could be linearly decoded both within and across recording sessions, we trained linear-kernel support vector machine classifiers (`sklearn.svm.SVC` with default *C* = 1.0 and one-vs-one multi-class strategy) to predict spatial bin identity from population activity. Each training sample consisted of the population activity vector at one spatial bin on one lap, drawn from the lapwise spatial tuning functions, with the corresponding bin index (1, …, 36) as the target label. For intra-session decoding, we used leave-one-lap-out cross-validation within each session. For inter-session decoding, we trained on all laps of one session and evaluated on each lap of another session, restricting both sessions to neurons that were successfully tracked across the pair. At test time, predicted bin indices were converted to two-dimensional spatial coordinates by mapping each bin to its centroid (the mean mouse position within that bin), and performance was quantified as the mean of the *R*^2^ scores for the x and y coordinates.

Within-session decoding performance was highly reliable across all mice (R^2^ = 0.921 ± 0.040 SD across mice), with individual mice showing consistent spatial decoding accuracy (R^2^ range: 0.838 to 0.976). Brain region analysis revealed that visual (VIS) and retrosplenial (RSP) areas provided the strongest decoding performance both individually (VIS: 0.895 ± 0.042 R^2^; RSP: 0.889 ± 0.066 R^2^) and in combination (VIS + RSP: 0.917 ± 0.049 R^2^), while somatosensory regions showed more variable performance (SSp: 0.706 ± 0.105 R^2^; MO: 0.639 ± 0.285 R^2^), which aligns with our spatial consistency results.

Cross-session decoding revealed characteristic temporal dynamics in representation stability. When decoding between consecutive sessions (Δt = 1), performance remained high across most session pairs (R^2^ = 0.864 ± 0.081 SD across mice). However, decoding performance gradually declined as the temporal separation between training and test sessions increased, with sessions separated by maximum intervals showing substantially reduced accuracy (**Supplementary Fig. 2**, R^2^ = 0.223 ± 0.185 when Δt = 13). Interestingly, individual mice exhibited distinct patterns, with decoding performance clustering into two groups (**Supplementary Fig. 2B**) that maintained consistent relative performance across session pairs.

### B.3 Preserved covariation of population embeddings under drift implies orthogonality

The finding that position-by-position cross-correlation matrices are preserved across sessions raises the question of whether the inter-session transformation must itself be orthogonal. We show that this is true on the representational subspace, provided population norms scale uniformly across sessions. This provides theoretical justification for the Procrustes analysis and explains why rotation-like dynamics are consistently observed in representational drift studies.

Let 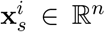 denote the population activity vector for position bin *i* ∈ {1, …, *m*} in session *s*, with *m* = 36 and *n* the number of neurons. Let 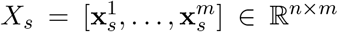 denote the matrix of population vectors, and let 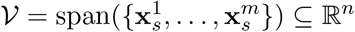 denote the representational subspace.

We make the following assumptions:

1. *Correlation preservation:* 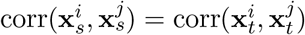 for all position bins *i, j*.
2. *Linear inter-session transformation:* there exists *R* ∈ ℝ^*n×n*^ such that 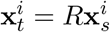 for all *i*.
3. *Uniform norm scaling:* 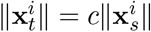 for all *i*, where *c >* 0 is constant.

Under these assumptions, the restriction of *R* to the representational subspace satisfies

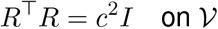

so *R* acts as an orthogonal transformation on 𝒱 up to the scale factor *c* (and is exactly orthogonal on 𝒱 when *c* = 1).

#### Proof

First, we express the correlation preservation condition where for any position bins *i* and *j*:

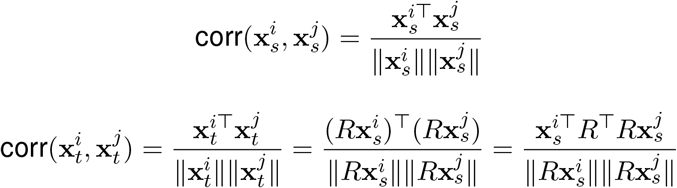

Then, by our first assumption

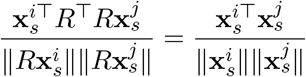

Next, we use the uniform norm scaling assumption that says that 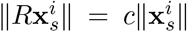 for all *i*. Therefore

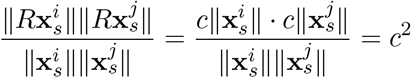

Substituting into the equation from the first step we get

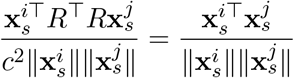

Therefore

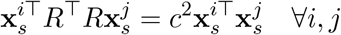

Which can be rewritten in matrix form as

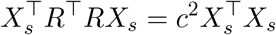

Finally, we need to show that *R*^⊤^*R* = *c*^2^*I* on the representational subspace.

To do so, note that for any vector **v** ∈ 𝒱, we can write **v** = *X*_*s*_***α*** for some ***α*** ∈ ℝ^*m*^. Then

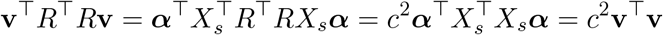

This shows that ∥*R***v**∥^2^ = *c*^2^∥**v**∥^2^ for all **v** ∈ 𝒱. By the polarization identity we have

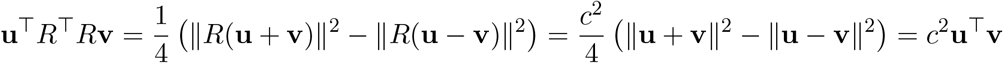

for all **u, v** ∈ 𝒱.

Therefore *R*^⊤^*R* = *c*^2^*I* on 𝒱.

## C Supplementary Figures

**Supplementary Figure 1:**
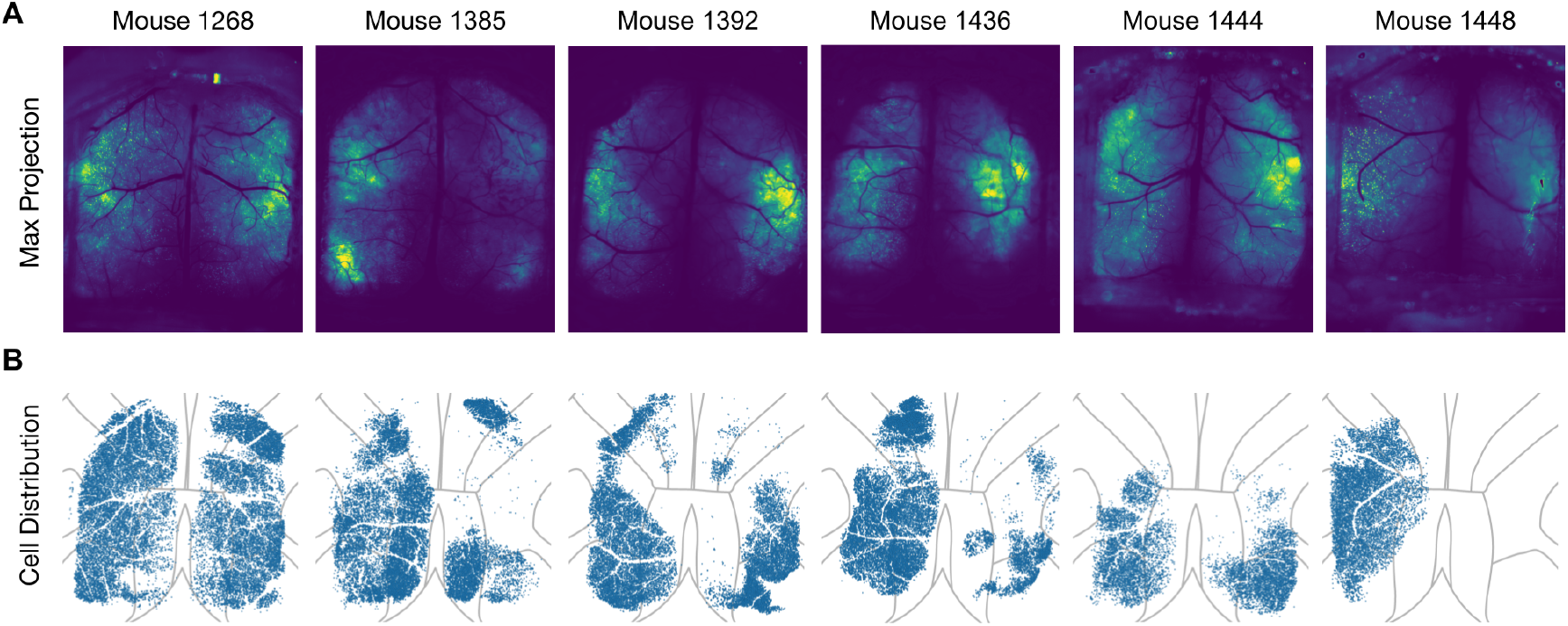
Imaging fields of view and anatomical distribution of tracked neurons across mice. **(A)** Maximum-intensity projections from session 1 of each mouse, produced by Suite2p. **(B)** Anatomical distribution of tracked neurons in each mouse, overlaid on the Allen Mouse Brain Atlas.

**Supplementary Figure 2:**
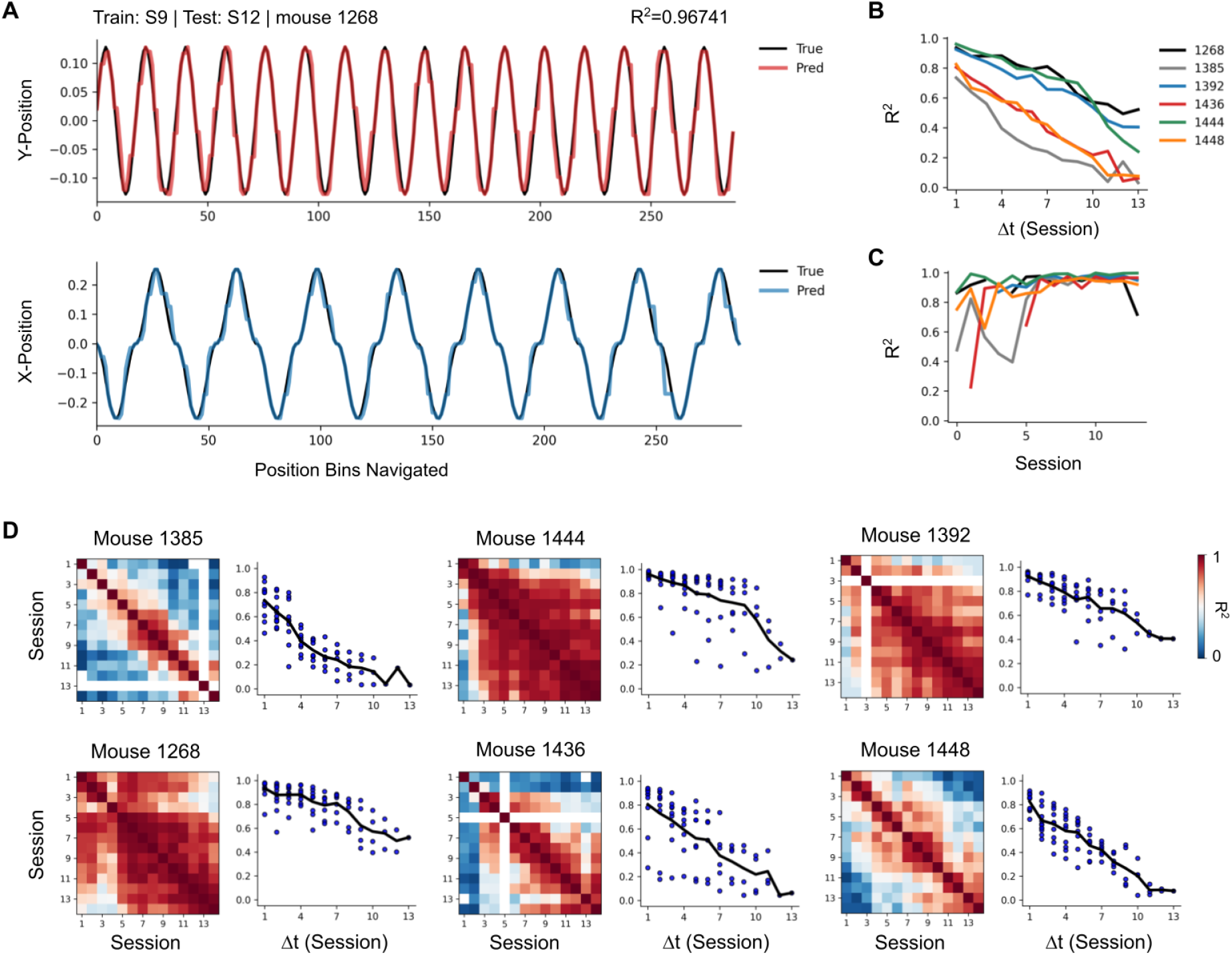
Linear decoding of mouse position from population activity within and across sessions. **(A)** Example cross-session decoding performance for X (blue) and Y (red) axes using mouse 1268, with a decoder trained on session 9 and evaluated on session 12. **(B)** Mean inter-session decoding performance versus session separation shown for each mouse. **(C)** Mean intra-session decoding performance within each session shown for each mouse. **(D)** Cross-session decoding performance matrices for each mouse. Dark red indicates high performance (R^2^=1), while dark blue indicates low performance (R^2^=0). Adjacent to each matrix is the decoding performance as a function of session separation.

**Supplementary Figure 3:**
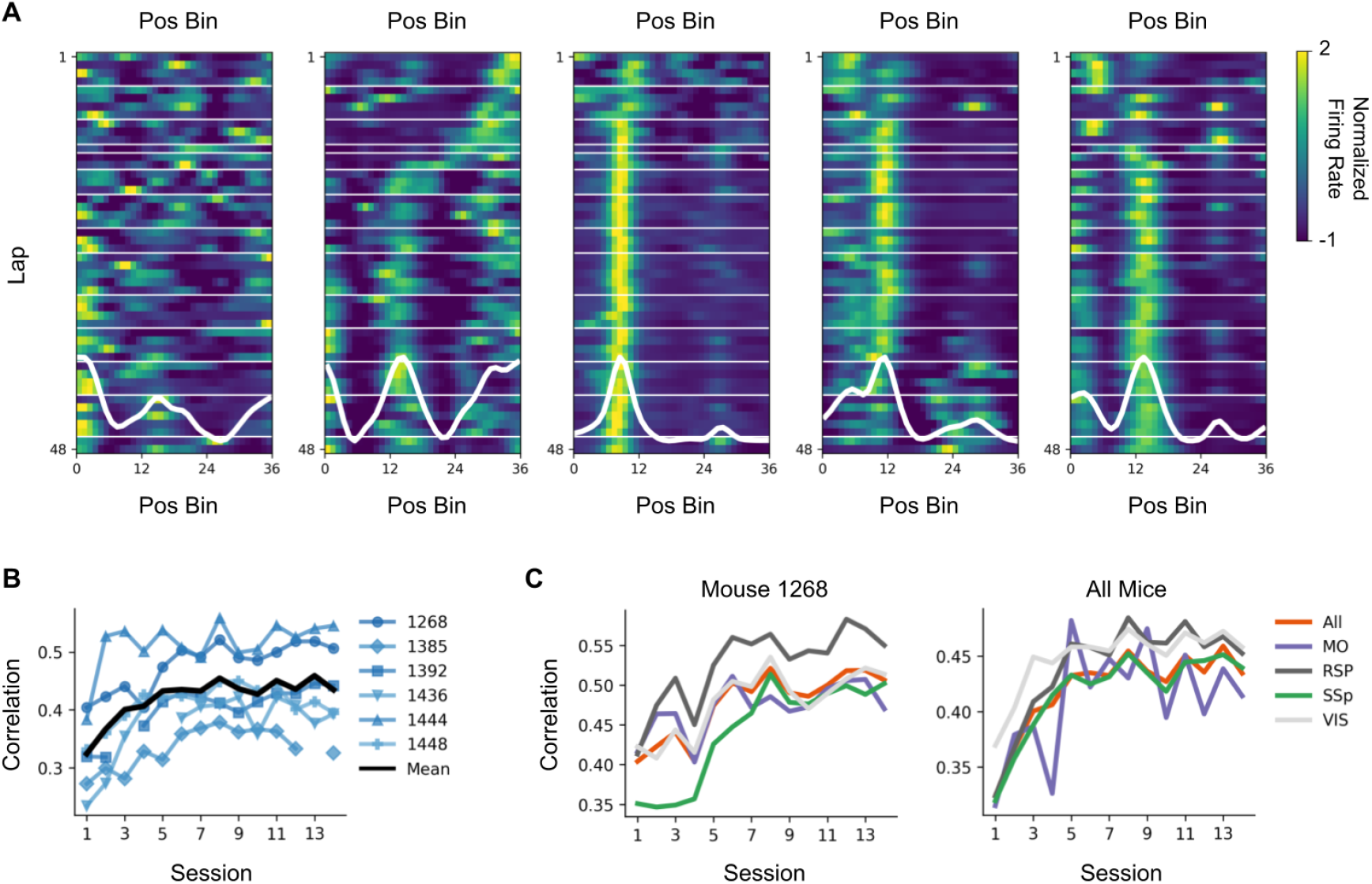
Spatial tuning consistency increases over the recording period across cortical regions. **(A)** Sample LSTFs from mouse 1268 that show this increase in consistency over time. **(B)** Correlation to the mean tuning averaged across all neurons for each mouse, then averaged across all mice shown in black. **(C)** The correlation to the mean tuning averaged across all neurons in each brain region for mouse 1268 (left) and all mice (right).

